# Determination of the structure and dynamics of the fuzzy coat of an amyloid fibril of IAPP using cryo-electron microscopy

**DOI:** 10.1101/2022.05.29.493873

**Authors:** Z. Faidon Brotzakis, Thomas Löhr, Steven Truong, Samuel E. Hoff, Massimiliano Bonomi, Michele Vendruscolo

## Abstract

In recent years, major advances in cryo-electron microscopy (cryo-EM) have enabled the routine determination of complex biomolecular structures at atomic resolution. An open challenge for this approach, however, concerns large systems that exhibit continuous dynamics. To address this problem, we developed the metadynamic electron-microscopy metainference (MEMMI) method, which incorporates metadynamics, an enhanced conformational sampling approach, into the metainference method of integrative structural biology. MEMMI enables the simultaneous determination of the structure and dynamics of large heterogeneous systems by combining cryo-EM density maps with prior information through molecular dynamics, while at the same time modelling the different sources of error. To illustrate the method, we apply it to elucidate the dynamics of an amyloid fibril of the islet amyloid polypeptide (IAPP). The resulting conformational ensemble provides an accurate description of the structural variability of the disordered region of the amyloid fibril, known as fuzzy coat. The conformational ensemble also reveals that in nearly half of the structural core of this amyloid fibril the side-chains exhibit liquid-like dynamics despite the presence of the highly ordered network backbone of hydrogen bonds characteristic of the cross-β structure of amyloid fibrils.

## Introduction

In the last several years, cryo-electron microscopy (cryo-EM) has been pushing the boundaries of structural biology in terms of structure resolution, system complexity, and macromolecular size^1,2^. Imaging single particles by rapid cryo-cooling and vitrification^3,4^ enables structural studies under near-native environmental conditions while offering sample protection from beam radiation. Technical advances in electron detectors, computational algorithms accounting for beam-induced motion, and automation of data collection and image analysis^5^ have paved the way to a spectacular increase of high-resolution cryo-EM density maps. The Electron Microscopy Data Bank (EMDB) currently holds 15,800 single-particle cryo-EM density maps, which offer exquisitely detailed structural information about macromolecular systems of central importance in cell biology.

In standard cryo-EM structure determination, two-dimensional (2D) images of single particles are first classified in conformationally homogenous classes and then averaged in a computational image processing step, thereby leading to a substantial increase in the signal-to-noise ratio, and thus in structure resolution^6,7^. However, the continuous dynamics of flexible regions are difficult to detect, therefore complicating the generation of homogenous classes of structures. The resulting low densities cannot be readily used to determine atomistic structures, and are thus often excluded from the final structural model. While methods such as the manifold embedding approach^8^ can determine multiple structures from cryo-EM density maps, to account for the conformational dynamics one should quantitatively and atomistically interpret the cryo-EM density maps as an envelope that corresponds to an averaged conformational ensemble of states with certain populations that interconvert with a characteristic timescale^9^. Such a viewpoint moves away from a single structure interpretation of the data and links the data to the statistical mechanics concept of free energy landscapes of conformational ensembles^10,11^. Integrative structural ensemble-modelling methods incorporate experimental information into molecular simulations and enable the determination of structural ensembles that maximally conform to the experimental data with atomistic resolution^12-23^. This technique has been applied using nuclear magnetic resonance (NMR) spectroscopy^17,19,20,24-28^, fluorescence resonance energy transfer (FRET) microscopy^29^, small-angle scattering techniques (SAXS/SANS)^30-34^, transition rate constants^22,23^ and cryo-EM data^35^.

One of such methods, cryo-EM metainference (EMMI)^35^, can accurately model a thermodynamic ensemble by combining prior information on the system, such as physico-chemical knowledge (e.g. a force field), with noisy (i.e. subject to systematic and random errors) and heterogeneous (i.e. encoding a conformational ensemble) experimental data, using cryo-EM density maps. EMMI has already been used in a series of complex macromolecular systems, including a CLP protease^36^, microtubules^37^, microtubule-tau complexes^38^, ASCT2 transporter^39^, and SARS-CoV-2 membrane receptor proteins^40^, allowing access to the continuous dynamics of biomolecules with atomistic resolution. The quality of the EMMI structural ensembles, however, is closely related to the exhaustiveness of the conformational sampling, which requires a computational time that scales exponentially with the barriers delimiting individual structural states^9^.

To tackle this rare event problem, several enhanced sampling methods have been developed. Enhanced sampling molecular simulation methods can be classified as trajectory based^41-46^ and collective-variable (CV) based^47-51^. For detailed reviews, readers can refer to the recent literature^52,53^. A particularly powerful CV-based enhanced sampling method, which is very efficient once appropriate CVs are chosen, is metadynamics^51,54^. Metadynamics adds a history-dependent bias to the system as a function of microscopic degrees of freedom of the system known as collective variables. With this bias, the simulations can escape deep free energy minima and sample transitions between different states. The choice of the CVs is critical to achieve the desired speed-up of convergence^52^. Recent developments in identifying and automating the search for appropriate CVs have increased the efficiency of this method, thus providing a remedy to the conformational sampling problem^55-59^. While metainference has been combined already with metadynamics^60^, this was not the case for EMMI (**Figure 1A**). Combining EMMI with enhanced sampling methods can lead to accurate and efficient determination of structural ensembles using the vast dataset of cryo-EM database, which can in turn provide atomistic insight to a range of biomolecular systems and processes.

**Figure 1.**
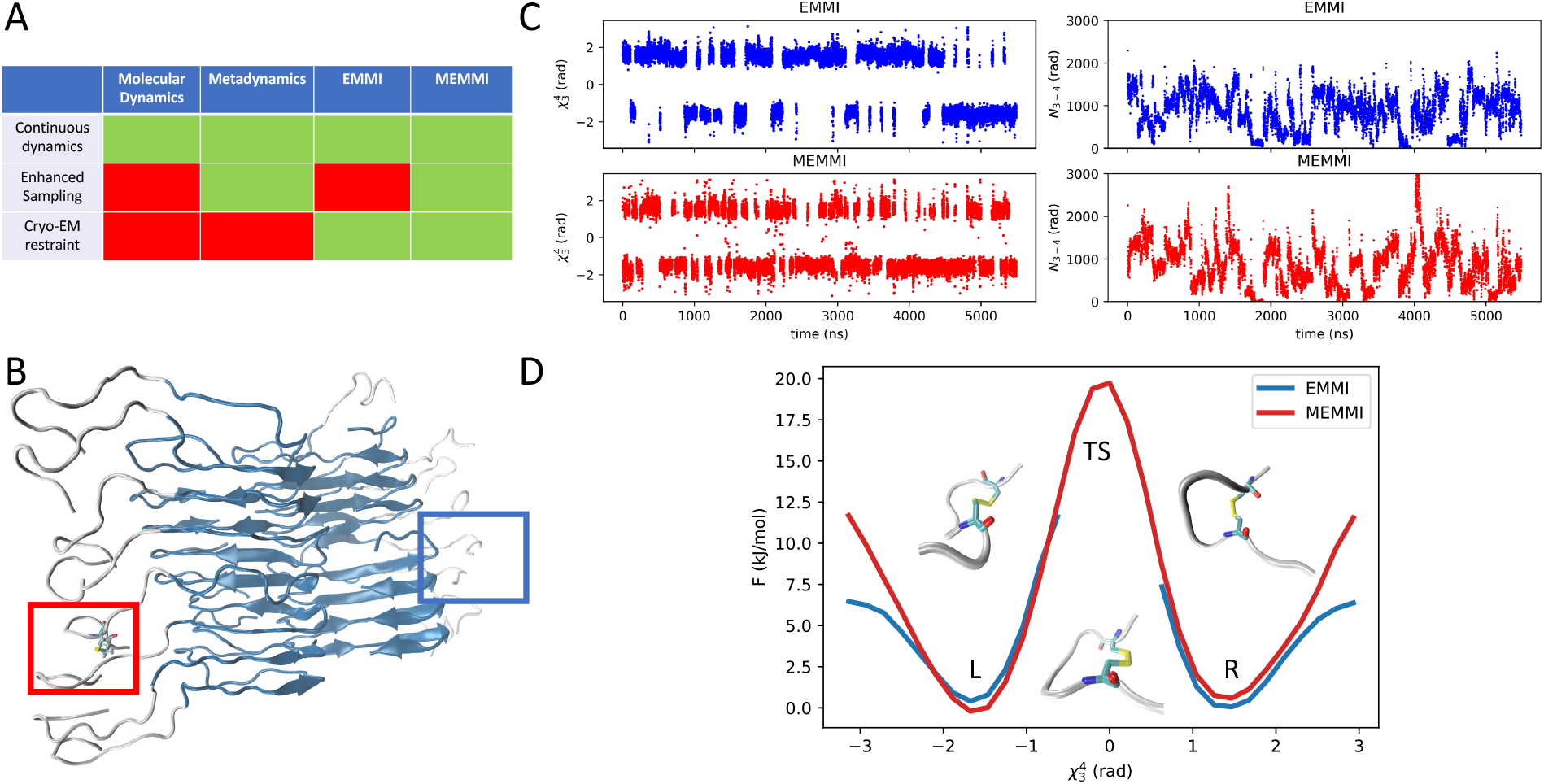
Acceleration of the conformational sampling using MEMMI. **(A)** Summary of the capability of the different approaches discussed in this work (molecular dynamics, metadynamics, EMMI and MEMMI) to include continuous dynamics, enhanced sampling and cryo-EM restraints. **(B)** A representative configuration of an IAPP amyloid fibril with a biased disulfide bond (red) and an unbiased one (blue). **(C)** Time trace of the χ_3_^4^ and N_3-4_ CVs with EMMI and MEMMI. **(D)** Free energy profile as a function of χ_3_^4^ with the corresponding structural states of the disulfide bond: MEMMI (red) EMMI (blue).

In this work, we present the MEMMI method, which incorporates metadynamics in EMMI in order to accelerate the ability of EMMI to sample structural ensembles with slowly interconverting states. We illustrate the application of this approach to determine the structural ensemble of an amyloid fibril formed by the full-length (residues 1-37) islet amyloid polypeptide (IAPP), an aberrant assembly associated with the degeneration of pancreatic β-cells in type-2 diabetes (T2D). When functioning correctly, IAPP, together with insulin, contributes to glycaemic control. IAPP and insulin are synthesized and stored together in pancreatic β-cells, but when IAPP aggregates in the extracellular space of the islets of Langerhans, amyloid-induced apoptosis of β-cells may occur^61^. In 95% of T2D patients, IAPP is found as extracellular amyloid deposits^62-64^, which form through surface-catalysed secondary nucleation^65^. IAPP fibrils represent a challenging system for structural biology studies, since no unique structure can be resolved in the low-density regions of the 12-residue long N-terminal tails, due to conformational heterogeneity and associated errors in the measurement. For this reason, although recent cryo-EM experiments determined various amyloid fibril structures of IAPP^66-68^, the structural heterogeneity in the disordered flanking regions, known as fuzzy coat, has so far proved impossible to resolve accurately. It would be desirable to acquire a better understanding of the conformational properties of the fuzzy coat, since this region is thought to play a central role in the interactions of amyloid fibrils with other cellular components such as RNA molecules and molecular chaperones^69,70^. Moreover, the fuzzy coat is likely to be involved in cell membrane binding, potentially promoting the catalysis of aggregation and capturing amyloid precursors^69,70^. Recent studies on the tau protein, which is implicated in numerous neurodegenerative diseases known as tauopathies, show that depending on pH conditions, the thick fuzzy coat changes the fibril properties, including mechanically stiffness, and repulsive and adhesive behaviours^69,70^. Here, we detail the continuous dynamics of the fuzzy coat of an IAPP fibril and utilise a thermodynamical theory of melting to characterize the different regions of the fibril to gain insight into its mechanical properties.

## The MEMMI method

### The cryo-EM forward model

A cryo-EM density map resulting from class-averaging and 3D reconstruction is typically encoded as voxels on a grid, and the map is generally distributed in this form. For computational efficiency, and to enable differentiability and analysis of correlations between data points, the map can be converted to a Gaussian mixture model *ϕ*_*D*_(***x***) (GMM) consisting of *N*_*D*_ Gaussian components

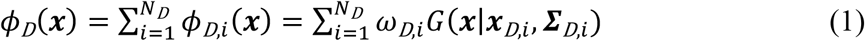

where ***x*** is a vector in Cartesian space, *ω*_*D,i*_ is the scaling factor of the i-th component of the data GMM and *G* is a normalized Gaussian function centered at ***x***_*D,i*_ with covariance matrix ***Σ***_***D***,***i***_. The agreement between models generated by molecular dynamics (MD) and the data GMM is calculated by the following overlap function *𝑜𝑣*_MD,*i*_

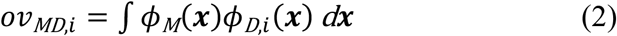

where *ϕ*_*M*_ (***x***) corresponds to the model GMM obtained from molecular dynamics. To deal with the heterogeneity of the system, EMMI simulates many replicas, *r*, of the system. The overlap between model GMM and data GMM is estimated over the ensemble of replicas as an average overlap per GMM component 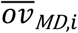. This forward model overlap can then be compared to the data GMM self-overlap *𝑜𝑣*_*DD,i*_ = ∫ *ϕ*_*D*_(***x***)*ϕ*_*D,i*_(***x***) *d****x***

### Metainference

Metainference is a Bayesian approach for modelling statistical ensembles by combining prior information on a system with experimental data subject to noise or systematic errors^17^. This framework is particularly well suited to structural ensemble determination through molecular dynamics simulations, in which the *prior* (i.e. the force field) is updated with information from experimental methods, such as NMR spectroscopy, SAXS, or cryo-EM data. Metainference is designed to handle systematic errors (such as biases in the force field or forward model), random errors (due to noise in experimental data), and errors due to the limited sample size of the ensemble^17^. The model generation is governed by the metainference energy function, defined as *E*_MI_ = −*k*_B_*T* log(*p*_MI_), in which *k*_B_ is the Boltzmann constant, *T* is the temperature, and *p*_MI_ is the metainference posterior probability

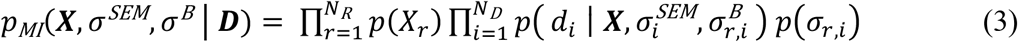

Here, **X** is a vector representing the atomic coordinates of the full ensemble, consisting of individual replicas *X*_*r*_; σ^SEM^ is the error incurred by the limited number of replicas in the ensemble; σ^B^ encodes the random and systematic errors in the prior, forward model, and experiment; and ***D*** = [*d*_*i*_] is the experimental data. Note *σ*^*SEM*^ is calculated per data point 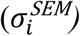, while *σ*^*B*^ is computed per datapoint *i* and replica *r* as 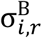. In the present case, the likelihood 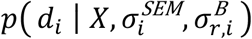 takes the form of a Gaussian function

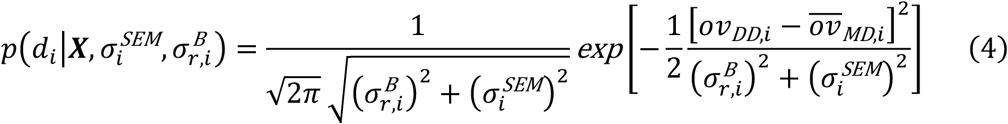

where 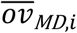 is the ensemble average of the overlap. The metainference energy function for multiple replicas then becomes

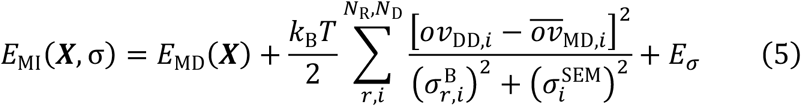

where *E*_*σ*_ represents the energy associated with the error *σ* = (*σ*^*B*^, *σ*^*SEM*^)

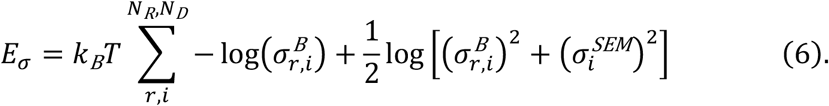

*E*_MD_ represents the molecular dynamics force field. While the space of conformations *X*_*r*_ is sampled by multi-replica molecular dynamics simulations, the error parameters for each datapoint 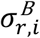 are sampled by a Monte Carlo sampling scheme at each time step. The error parameter related to the limited number of replicas used to estimate the forward model (*σ*^*SEM*^) can be chosen as a constant or estimated on the fly by using a windowed average^32^.

### Metadynamic cryo-EM metainference (MEMMI)

To accelerate the sampling of the metainference ensemble, one can utilise an enhanced-sampling scheme such as metadynamics^51,60^. In this case, we use parallel-bias metadynamics (PBMetaD)^55^ with the multiple walkers scheme^71^. Here, *V*_PB_ is a time-dependent biasing potential acting on a set of *N*_CV_ collective variables *s*(***X***), which in turn are functions of the system coordinates

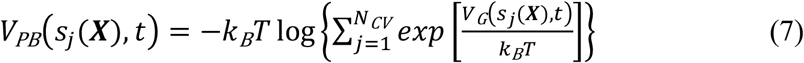

In contrast to conventional metadynamics, in PBMetaD multiple one-dimensional bias potentials *V*_G_ are deposited rather than a single high-dimensional one. This alleviates the curse of dimensionality while still allowing an efficient exploration of phase space^55^. Additionally, the use of multiple replicas through the multiple walkers scheme^71^ allows the sharing of the bias potential to drastically improve the sampling performance, while at the same time being a natural fit for the replica averaging approach of EMMI. Analogously to well-tempered metadynamics, these bias potentials *V*_G_(*s*_*j*_ (***X***), *t*) eventually converge to the free energy *F*(*s*_*j*_ (***X***)). The MEMMI energy function then becomes

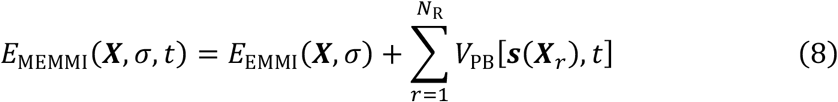

While the PBMetaD bias potential is shared among replicas, each replica may still experience a varying potential depending on its location in phase space. Thus, the arithmetic average over the forward models (i.e., the overlap) no longer presents an unbiased estimate of the ensemble average. It therefore needs to be replaced with a weighted average, utilising the bias potential of each replica *r* to unbias the ensemble at time *t*

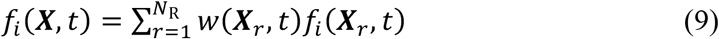

where the unbiasing weight *w*(***X***_*r*_, *t*) is defined as

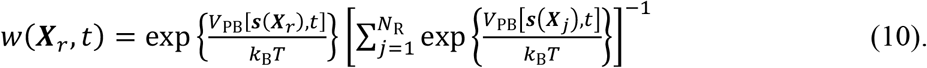

The unbiasing procedure used here is analogous to the standard umbrella-sampling technique by Torrie and Valleau^47^. Now the ensemble average 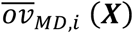 is given by

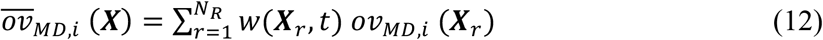

The MEMMI energy *E*_MEMMI_ is thus equal to Eq. 5.

### Initial fibril structure

We build the initial structure of the full fibril by starting from a deposited fibril structure (PDB:6Y1A), which only contains the core of the fibril, and extending each of the 16 peptide chains by adding the missing 12-residue N-terminal sequence with guidance from the cryo-EM density map EMD-10669. We use the macromolecular model-building program coot^72^.

### Molecular dynamics setup and equilibration

We continue by creating a 12.34 × 12.34 × 12.34 nm cubic simulation box, solvating with 58291 water molecules and neutralizing the net charge by adding 48 Cl^-^ ions. We use the CHARMM22*^73^ force field and TIP3P^74^ water models. We continue with an energy minimization followed by a 500 ps NPT equilibration at a temperature of 310 K and pressure of 1 atm, followed by an additional 2 ns NVT equilibration at 310 K. The molecular dynamics parameters are the same used previously^40^.

### MEMMI simulations

We first express the experimental voxel map data as a data GMM containing 10,000 Gaussians in total, resulting in a 0.975 correlation to the original voxel map EMD-10669^61^, using the gmmconvert utility^75^. We continue by extracting 32 configurations from the previous NVT equilibration and initiate a MEMMI simulation, consisting of 32 replicas, resulting in an aggregate runtime of 5.49 μs, using PLUMED.2.6.0-dev^76^ and gromacs-2020.6^77^. The simulation is performed in the NVT ensemble at 310 K using the same MD parameters as in the equilibration step. Configurations are saved every 10 ps for post-processing. The cryo-EM restraint is updated every 2 MD steps, using neighbour lists to compute the overlaps between model and data GMMs, with a neighbour list cut-off of 0.01 and update frequency stride of 100 steps. The biasing collective variables *s* = [*s*_*i*_] in the simulation are shown in **Figure S1A** and the biasing scheme is PBMetaD^55^ with the well-tempered^78^ and multiple-walkers^71^ protocols. The hill height is set to 0.3 kJ/mol, with a deposition frequency of 200 steps and an adaptive Gaussians diffusion scheme^79^. The biasing collective variables correspond to degrees of freedom of the left hand-side N-terminal. The respective degrees of freedom of the right hand-side N-terminal do not feel a metadynamics potential and therefore in the remaining text will be referred to as EMMI degrees of freedom and are also listed in **Figure S1**. As a post-processing step, we generate the final structural ensemble by resampling the generated configurations based on the converged unbiasing weights for each structure after an equilibration of 7 ns shown in **Figure. 1A**. To establish convergence, we perform a clustering analysis on the structural ensemble based separately on the first and second half of each replica (taking into accounts the weights) using the GROMOS method^80^, and with metric the root mean square deviation (RMSD) calculated on the Cα (CA) atoms (**Figure S1B**). Time-traces of CVs as well as their time-dependent free energy profile are shown in Figs. S2-S4. For molecular visualizations and calculating the local correlation of the final structural-ensemble-generated cryo-EM map with the experimental cryo-EM map, we use Chimera and gmconvert^81,75^. Except otherwise mentioned, all the structural analysis is performed on degrees of freedom of left side tail N-tail and chains 3 and 4. We do so in order to avoid the finite size effects on the outermost N-tails.

### Structure and dynamics of IAPP amyloid fibrils

#### Acceleration of the conformational sampling

MEMMI accelerates the conformational sampling by biasing a set of microscopic degrees of freedom of the system, also known as collective variables (CVs, **Figure S1)**. To demonstrate the performance of this approach, we visualize the time-trace of the collective variable χ_3_^4^ which corresponds to the disulfide bond dihedral of the 4^th^ peptide in the eight-layer stack of the fibril. This peptide is thus representative of a buried monomer with little interaction with the fibril ends. Note that, due to C2 helical symmetry, there are two of these dihedrals, one corresponding to the one side and one to the other sides N-terminal tail (**Figure 1B**). The sampling of one side is accelerated by a biasing potential of Eq. 7 (MEMMI), while the other is not (EMMI). Compared to the dihedral, the biased disulfide χ_3_^4^ (MEMMI) shows an increased transition rate, and thus more efficient conformational sampling (**Figure 1C)**. While both methods characterise the wells of the stable states (L, R), MEMMI is able to provide access to higher free energy transition state (TS) regions (**Figure 1D)**. Monitoring a non-periodic CV, such as the number of contacts between N-tail three and four, shows diffusion along high-low contact regions in both MEMMI and EMMI cases but somewhat more frequent in the MEMMI case. The combination of diffusion shown in the traces of CVs and the free energy profiles as a function of simulation time of all collective variables biased by EMMI and MEMMI are shown in **Figures S2-S4**, and indicate that our simulations are well converged. Taken together, these results show that both the MEMMI and EMMI simulations are converged in the low free energy regions, while MEMMI enables visiting high free energy regions.

#### Conformational heterogeneity of the fuzzy coat

A structure of the IAPP fibril core (residues 13-37) has been previously published (PDB:6Y1A) using the cryo-EM data used also in the present work (EMD-10669). Here, we determine a structural ensemble of the whole IAPP fibril (residues 1-37). We model the system as a stack of eight peptides per side (**Figure 2A**). While the core of the fibril maintains a parallel β-sheet structure, the flanking region (residues 1-12) exhibits a large conformational heterogeneity (**Figure 2B**). While we find the cores residues 12-37 to be largely in a β-sheet conformation, we also note significant heterogeneity for residues 23-24, 32-34 and 37. As shown in **Figure 2B**, residues 23-24 interact with TYR37 and maintain mostly a coil structure, while residues 32-34 interact with the fuzzy coat and maintain mostly a coiled structure. We also detected a small population of α-helical conformations in the region of residues 5-9.

**Figure 2.**
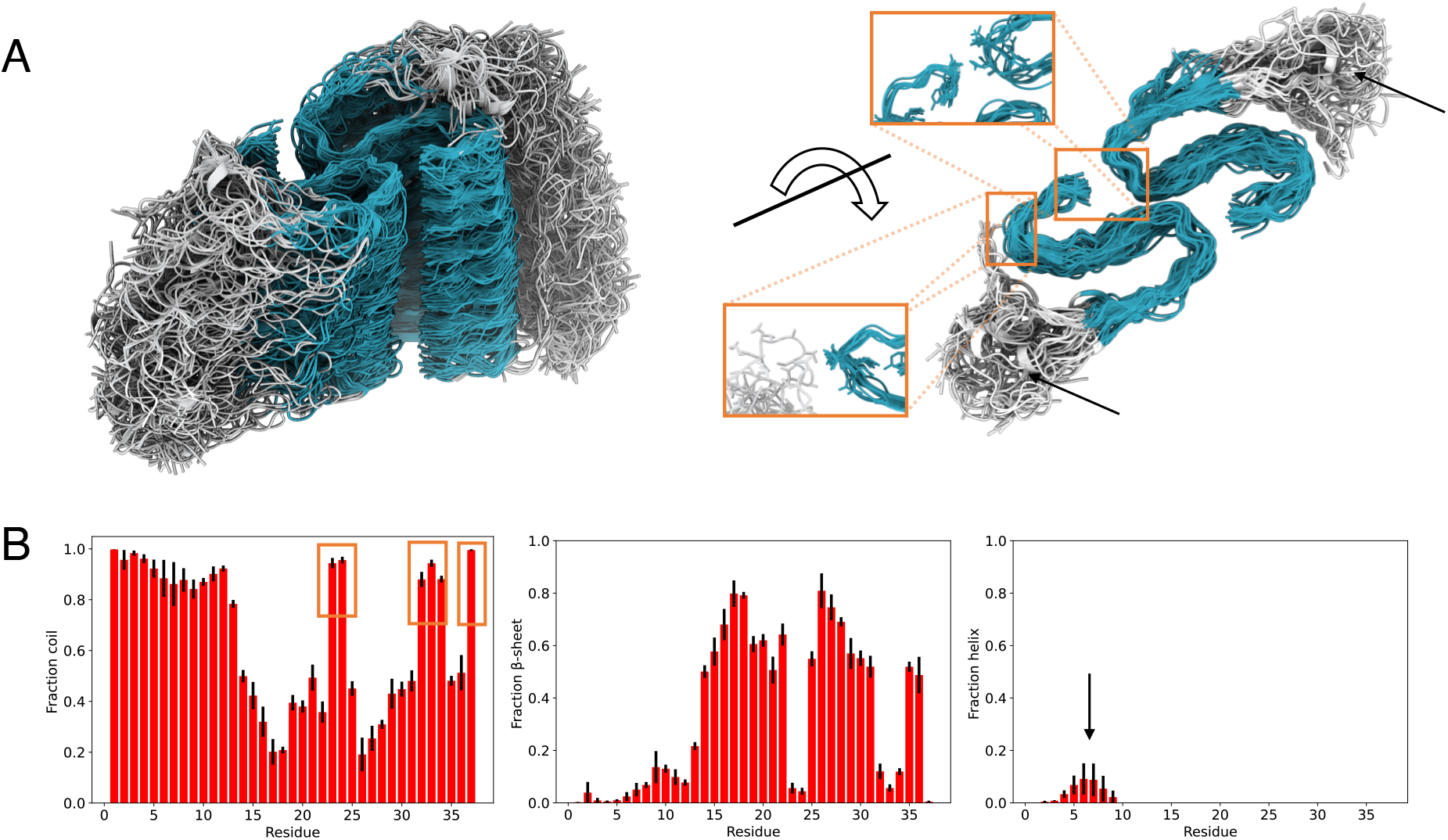
A structural ensemble of a IAPP amyloid fibril. **(A)** Core fibril residues (13-37) are shown in cyan, while the N-terminal tail residues (1-12) are shown in grey (fuzzy coat). This representation of the structural ensemble was generated by extracting 50 conformations from the final structural ensemble. Close-ups of interfaces are shown from 10 conformations. Examples of individual conformations are shown in **Figure S6. (B)** Structural analysis reporting on the fraction of coil, β-sheet and α-helix formed per residue in the ensemble obtained from MEMMI. Error bars are calculated as standard deviation between the first and second simulation half.

#### Correlation between experimental and calculated cryo-EM maps

We estimate the correlation of the MEMMI structural ensemble with the experimental cryo-EM map (**Figure 3**). We find that using a structural ensemble IAPP (residues 1-37) correlates better with the experimental cryo-EM map than a single structure (PDB:6Y1A) (**Figure 3A,B)**. The coefficient of correlation of the structural ensemble to the experimental electron density map is on average 0.92. Furthermore, an important feature of MEMMI is its ability to estimate the error in the experimental electron density map (**Figures 3C** and **S5**). We find that the relative error per Gaussian data point is on average 0.09, where the relative error is the error of each Gaussian data point with respect to the total overlap between all data GMM and the i^th^ component of the data GMM^35^. The low-density, high-error volume around residues S34, N35, and T36 can likely be attributed to the lack of MES/NaOH buffer in our MEMMI simulations, which is present in the experimental setup. Both MEMMI and EMMI exhibit good corelation (about 0.83) in the respective N-tail region with the cryo-EM map (**Figure 3D**).

**Figure 3.**
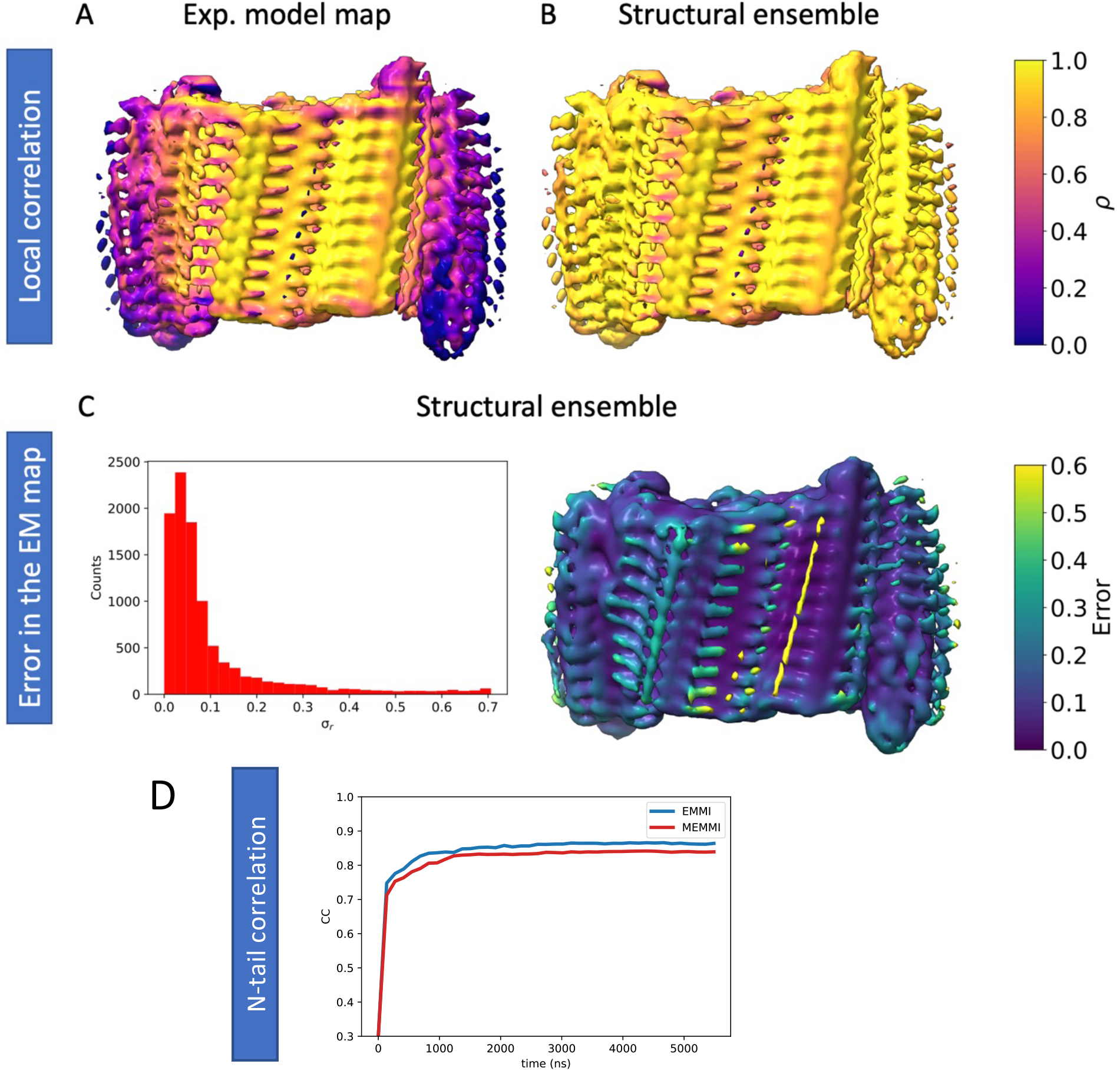
Local correlation and error in the data from MEMMI. **(A, B)** Assessment of the local correlation between the cryo-EM map (EMD-EM-10669) with cryo-EM maps generated by a previously determined single structure (PDB:6Y1A) **(A)**, and the MEMMI structural ensemble **(B). (C)** Histogram of the error in the GMM data as obtained by the structural ensemble (left). Error in the data, projected in the EMD-EM-10669 (right). **(D)** Time-dependent correlation of the left/right side N-tail structural ensemble (MEMMI/EMMI) with the corresponding region in the cryo-EM map.

#### Comparison of the dynamical properties of the fibril core and fuzzy coat

The conformational properties of the fuzzy coat and core region have been shown to be relevant modulating the properties of amyloid fibrils, including their ability to interact with various cellular components^69,70^. To investigate this phenomenon in the case of the present IAPP amyloid fibril, we take inspiration from a thermodynamic theory of melting and characterize the residue-dependent Lindemann parameter Δ_L_ (**Figure 4**), which encodes information on solid-like and liquid-like behaviour^82^. At the backbone-level, we find that the fibril core (23-30) is solid-like (Δ_L_ < 0.15), while the flanking region (1-12) is on the verge of a liquid-like behaviour (Δ_L_ ≥ 0.15). The Lindemann parameters of the side-chains indicate more mobility and are liquid-like outside the region of residues 20-32. These results reveal that about half of the structural core of this amyloid fibril remains rather disordered at the side-chain level, a phenomenon observed also for folded native states^82^. This is in contrast to monomeric disordered proteins such as amyloid-β, which generally exhibit Lindemann parameters Δ_L_ ≥ 1.0 (**Figure S7**).

**Figure 4.**
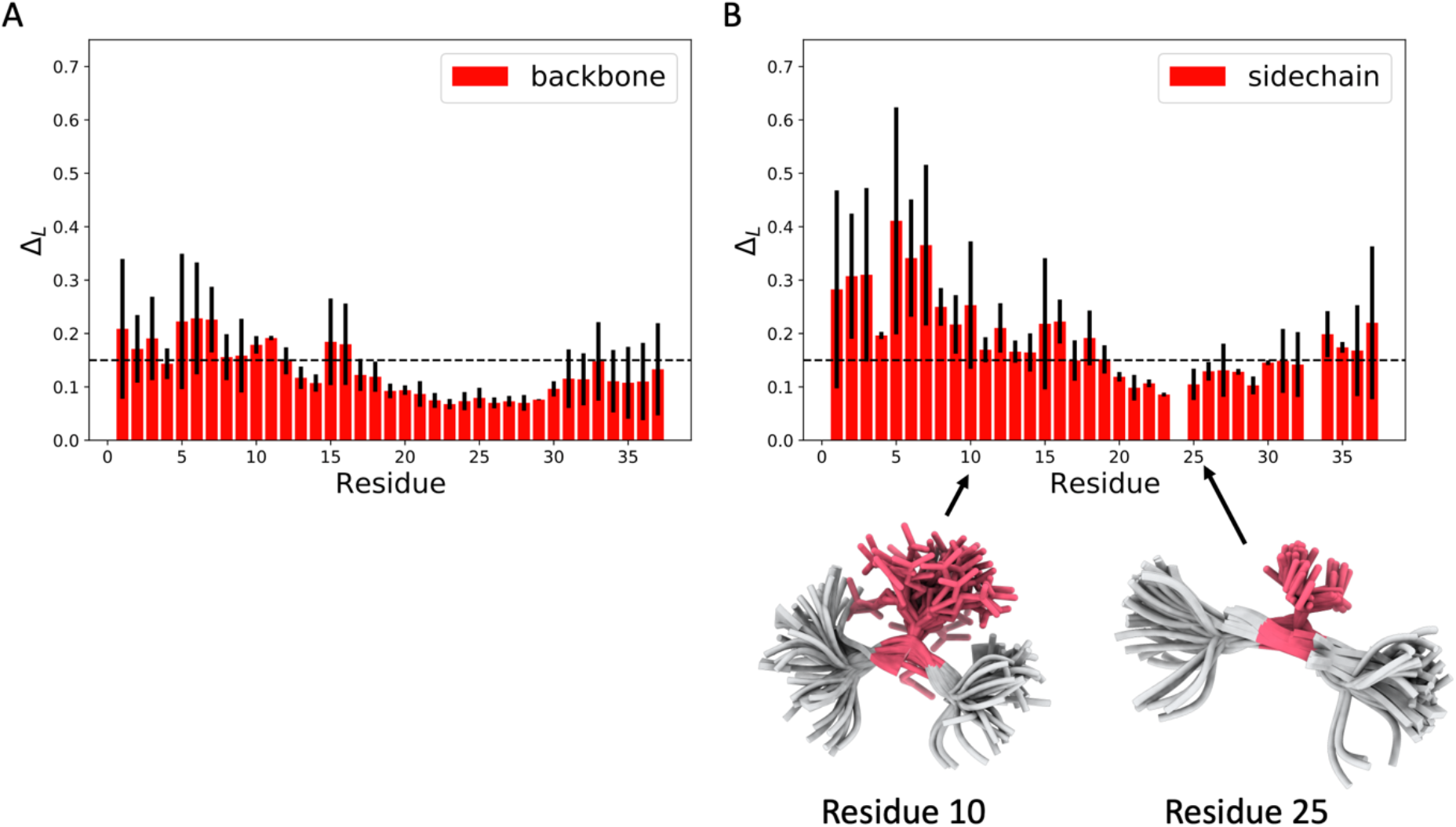
Analysis of the liquid-like and solid-like behaviours of the backbone and side-chains. **(A,B)** Per-residue Lindemann parameter (Δ_L_) for backbone **(A)** and side-chain **(B)** atoms. The dashed line at Δ_L_ = 0.15 marks the threshold value of the Lindemann parameter that distinguishes the solid-like and liquid-like behaviours. The behaviour of the side-chains is solid-like only in the region of residues 20-32, which is about half of the fibril core (residue 13-37). Error bars are calculated as standard deviation between the first and second simulation half.

#### Residue-specific solubility of the amyloid fibril

To investigate whether or not the surface of the amyloid fibril is soluble, we calculate the solubility per residue, using the structure-corrected CamSol solubility score^83^ (**Figure 5A,B**). We find three insoluble regions: (i) residues 5-7 (ATC) in the fuzzy coat, (ii) residues 26-29 (ILSS) at the ends of the fibril fragment, and (iii) residues 14-18 (NFLVH) on the fibril surface, stretched along the helical axis. The latter result is consistent with experimental evidence of residues N14, H18, and S20 being implicated in IAPP aggregation^63,84^ via secondary nucleation. We note that the solvent-exposed and aggregation-related residues 5-7 and 18-20 might present attractive targets for structure-based drug discovery.

**Figure 5.**
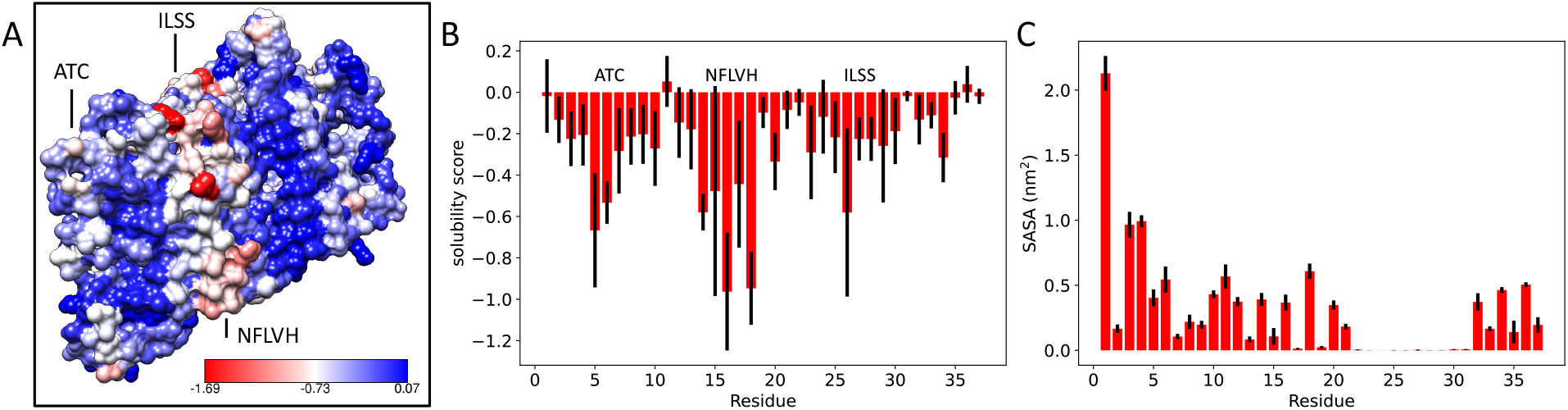
Analysis of the solubility of the fibril surface from the MEMMI ensemble. **(A)** Representative structure with CamSol solubility scores indicated on the surface (right). **(B)** Per-residue CamSol solubility score. **(C)** Per-residue solvent-accessible surface area. Error bars are calculated as standard deviation between the first and second simulation half.

## Conclusions

We have presented the MEMMI method for the simultaneous determination of the structure and dynamics of large and conformationally heterogeneous biomolecular structures from cryo-EM measurements. To illustrate the type of information that can be extracted from this type of approach, we have reported a structural ensemble of an amyloid fibril formed by IAPP. The analysis of the structural ensembles has revealed the conformational and the aggregation-related solubility properties of the fuzzy coat of the amyloid fibril, and that many of the side-chains in the structural core of the amyloid fibril exhibit liquid-like behaviour. Since this phenomenon has also been observed for native states of proteins, these results reveal a similarity in the structural behaviour of proteins upon folding in their native and amyloid states.

## Data availability

Input files for the simulations can be found at PLUMED-NEST (<u>plumID:22.023</u>) The ensemble can be found at https://zenodo.org/record/6518554#.YnLwby8Rqhw

## Acknowledgements

Z.F.B. would like to acknowledge the Federation of European Biochemical Societies (FEBS) for financial support (LTF). M.B. and S.E.H. acknowledge the support of the French Agence Nationale de la Recherche (ANR), under grant ANR-20-CE45-0002 (project EMMI).

## Supplementary Information

**Figure S1.**
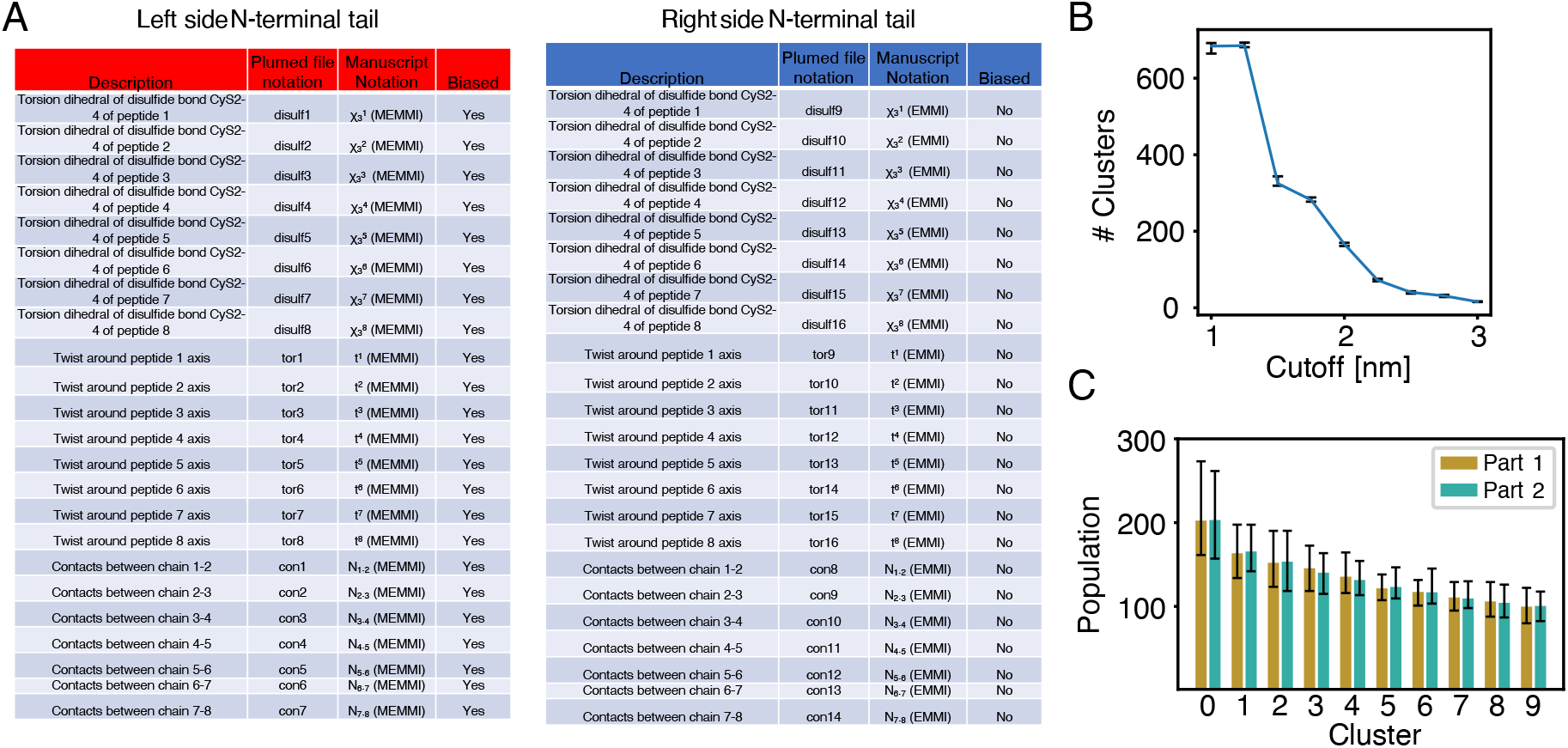
Metadynamics and clustering information. **(A)** Summary of all biased collective variables of the left side N-terminal tail and the respective unbiased ones on the right side N-terminal tail used in the analysis. **(B)** Dependence of number of clusters on the cut-off value used in the GROMOS clustering algorithm, using root-mean-square deviations of Cα atoms in tails 3 and 4 of the left (biased) side. **(C)** Populations of the top 10 clusters for the two last 40 % chunks of the simulation. Error bars for **(B)** and **(C)** show the 95th percentiles over 20 separate clustering runs using 5000 frames sampled based on metadynamics weights from each part of the simulation.

**Figure S2.**
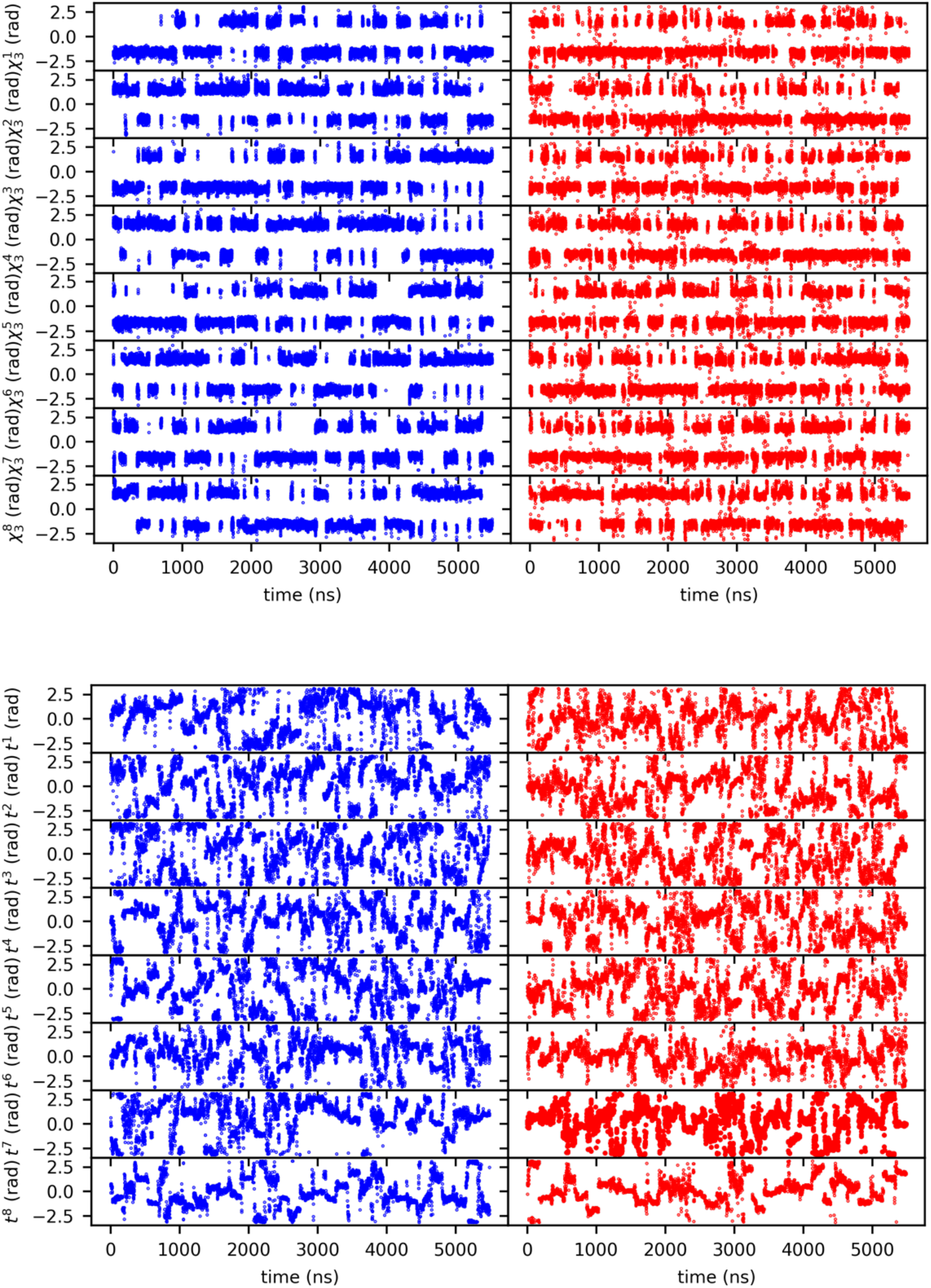
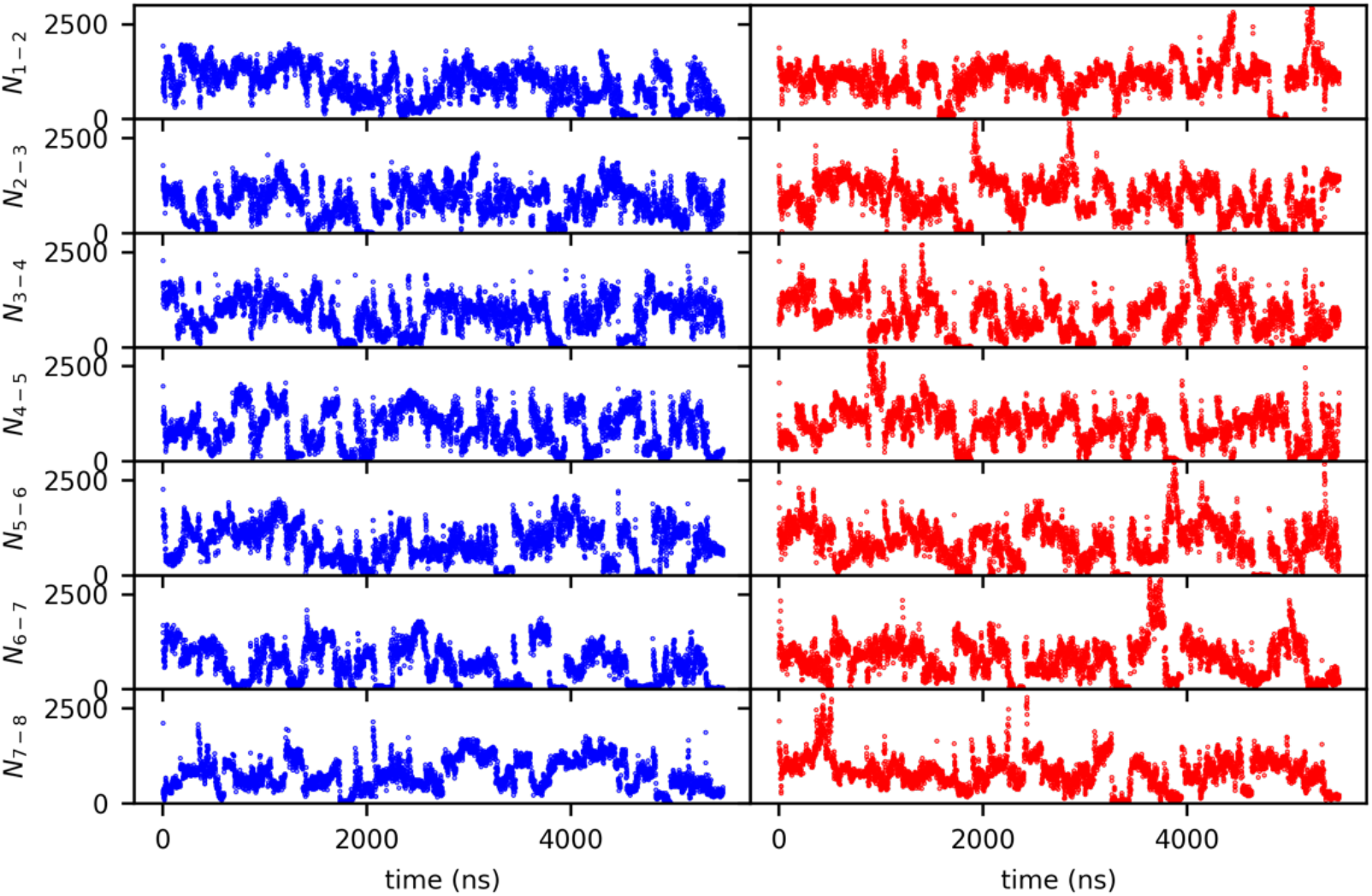
Assessment of the convergence of the simulations MEMMI-EMMI. Time evolution profiles for all unbiased (blue) and respective biased CVs (red).

**Figure S3.**
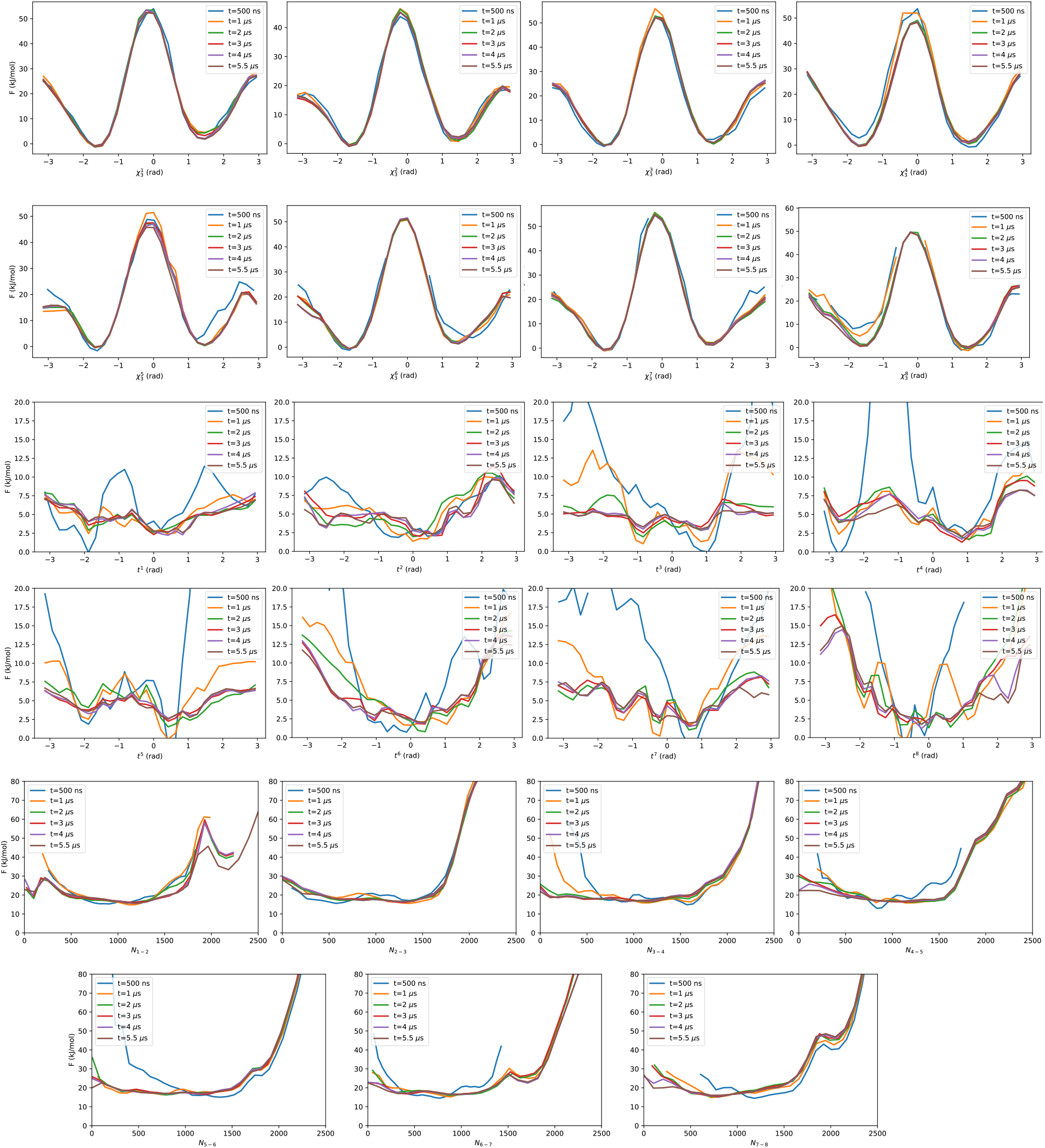
Assessment of the convergence of the MEMMI simulation. Free energy profiles for all biased collective variables for subsequent 1 μs increments of simulated time.

**Figure S4.**
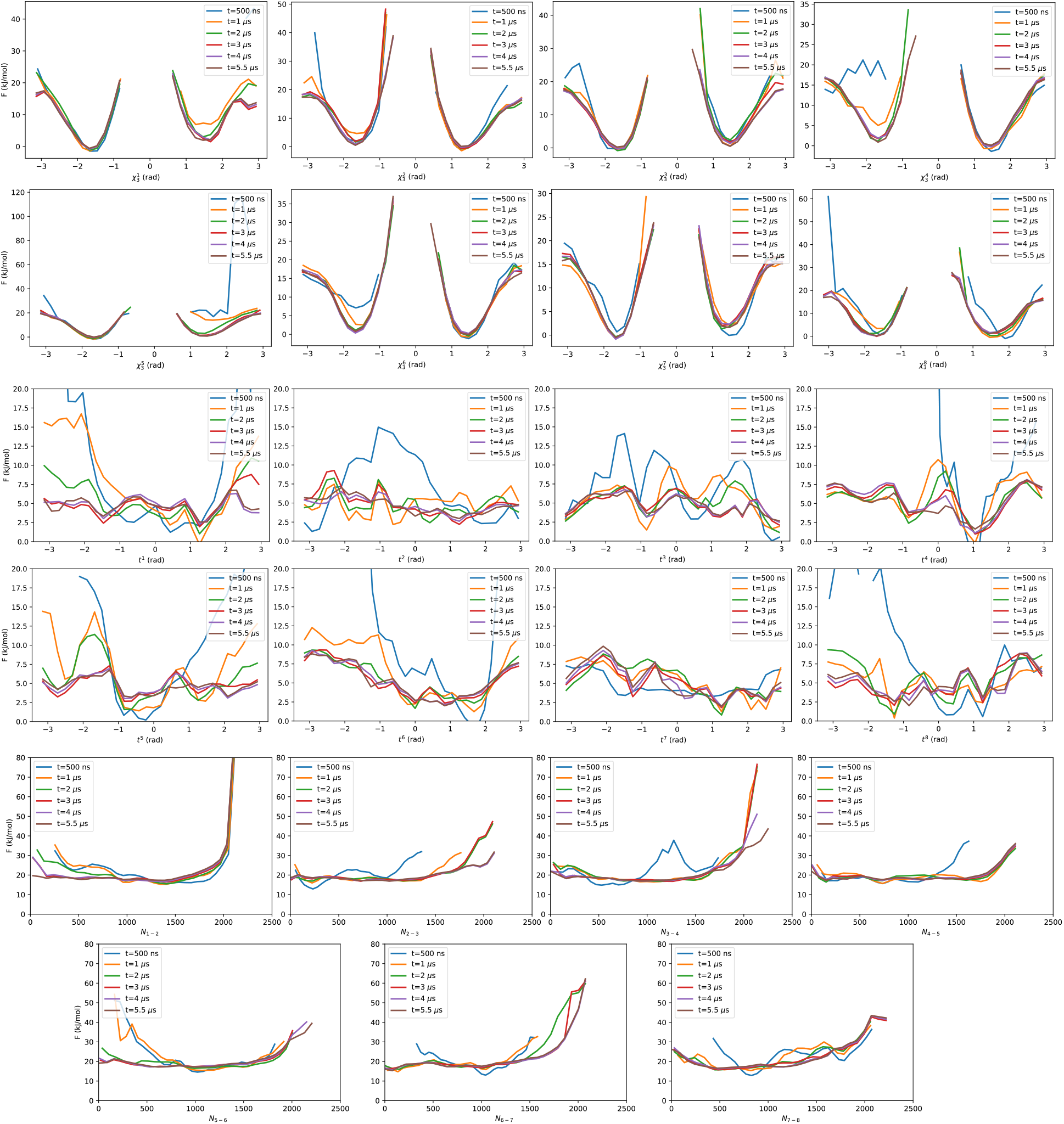
Assessment of the convergence of the EMMI simulation. Free energy profiles for all unbiased collective variables for subsequent 1 μs increments of simulated time.

**Figure S5.**
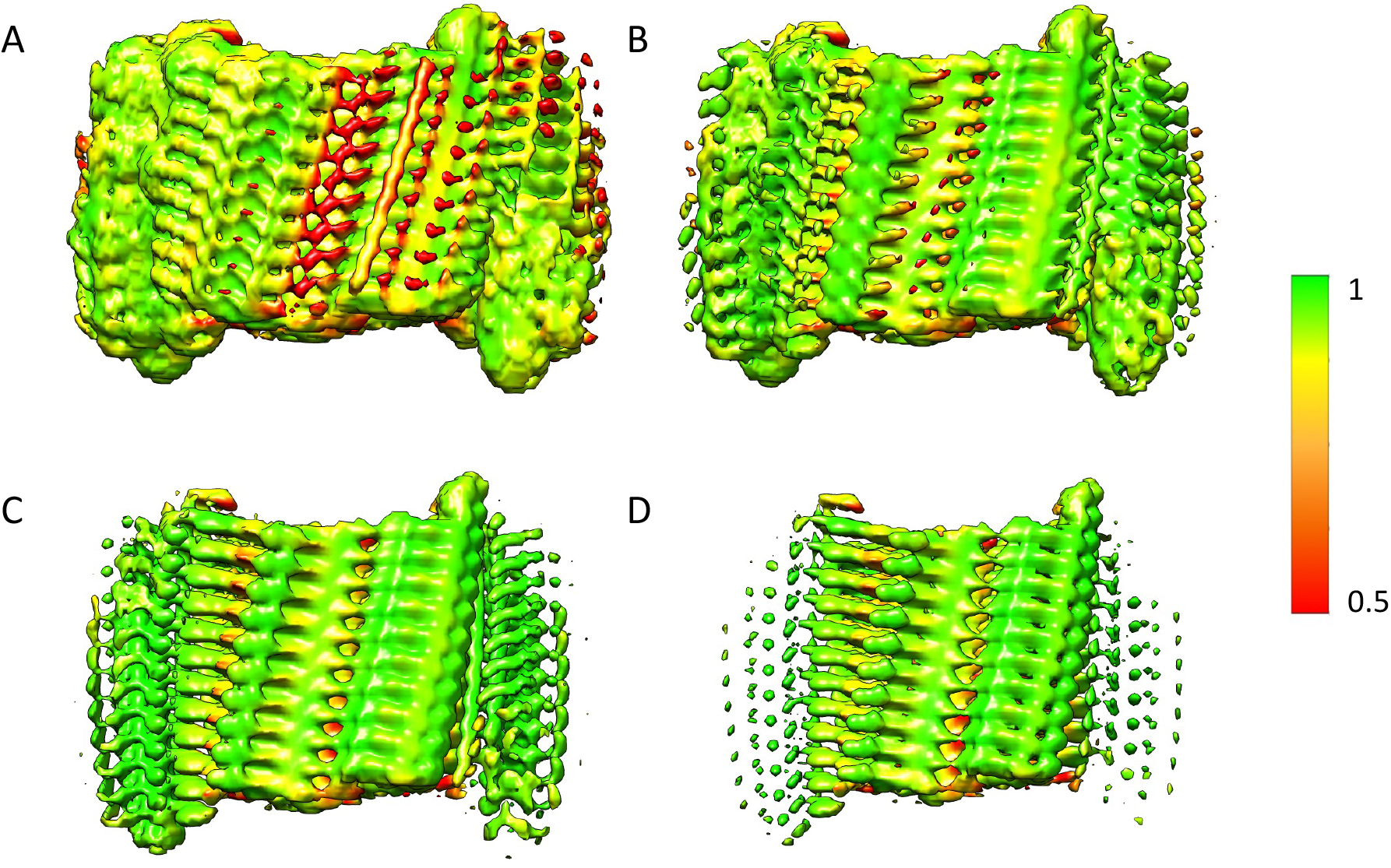
Assessment of the local correlation between experimental and calculated cryo-EM map. Local correlation of the cryo-EM map (EMD-EM-10669) with a map generated from the MEMMI ensemble of an IAPP amyloid fibril as a function of increasing strength of density (decreasing electron density thresholds): 1σ (A), 2σ (B), 3σ (C), and 4σ (D).

**Figure S6.**
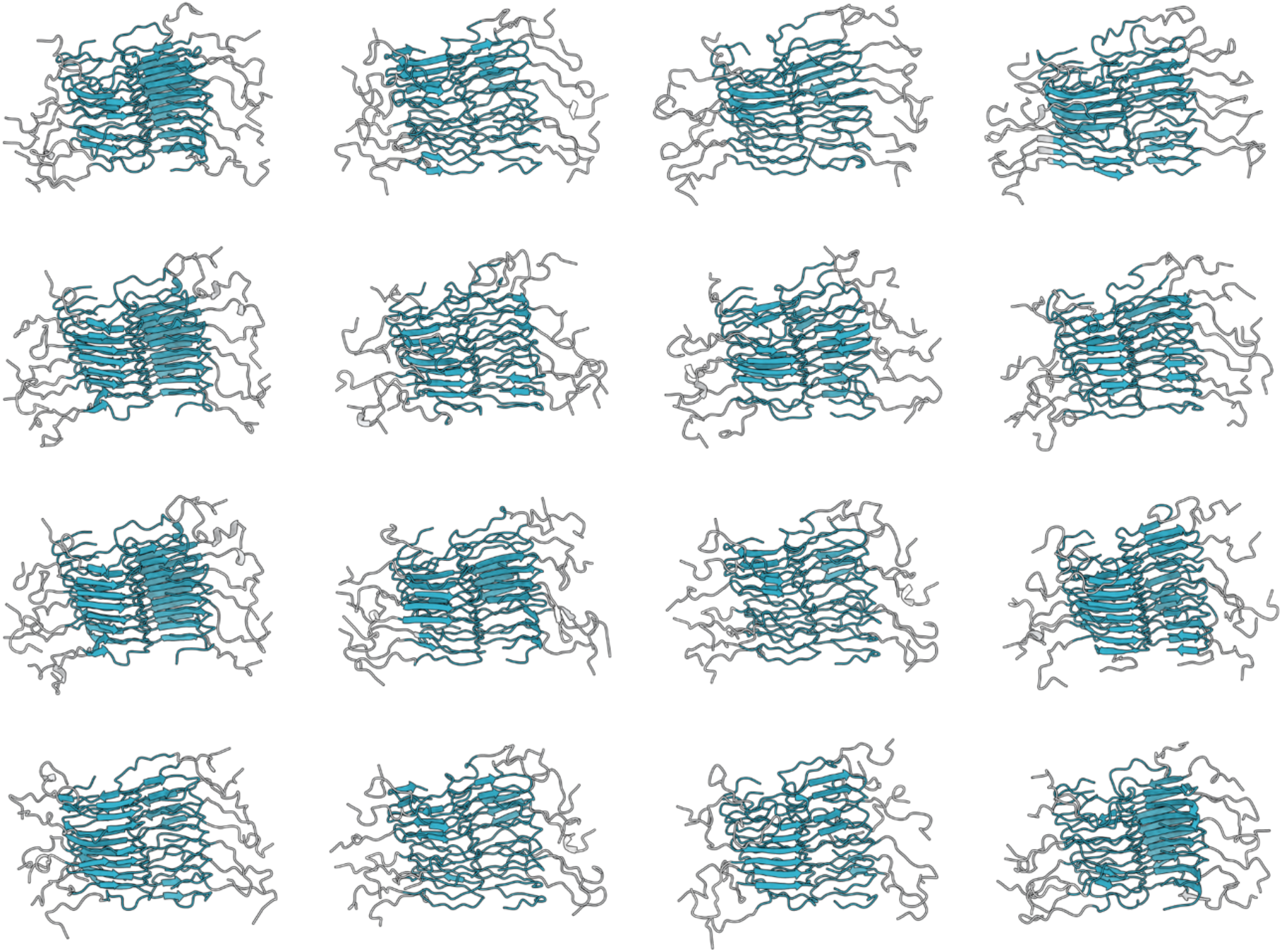
Individual conformations of the final structural ensemble. Sample of 16 structures from the ensemble shown in **Figure 2**.

**Figure S7.**
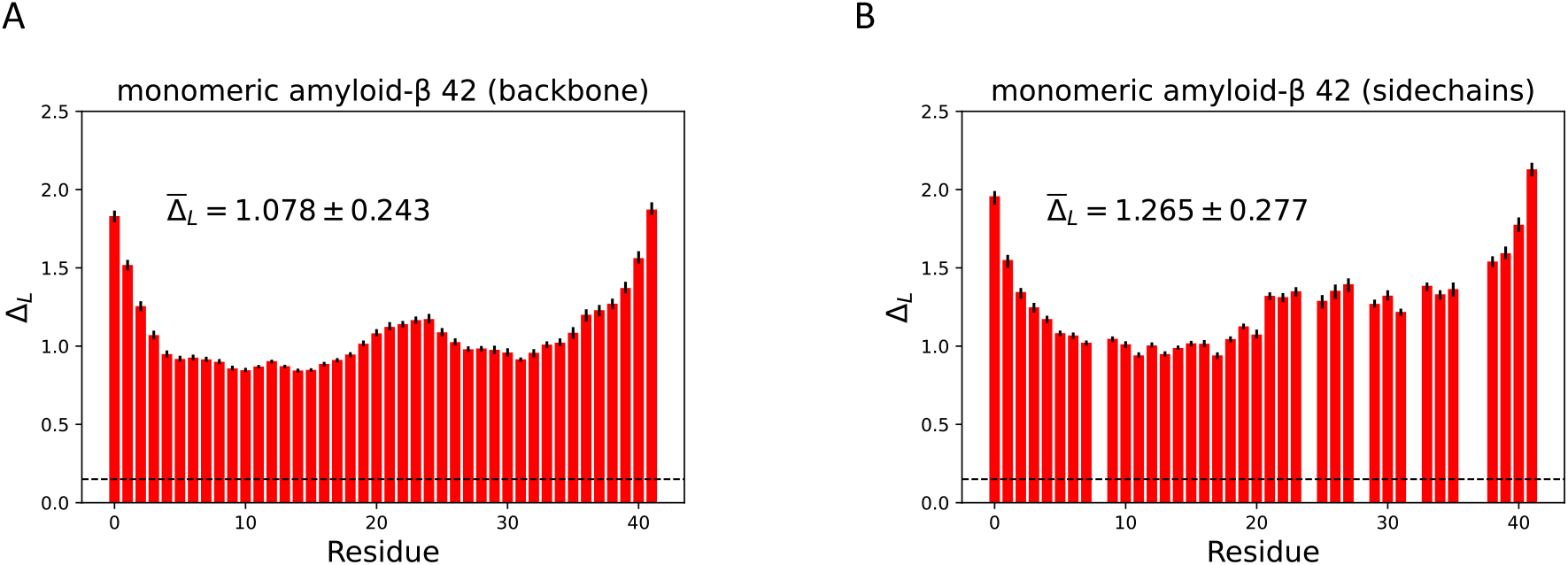
Lindemann parameters for the disordered monomeric amyloid-β 42 peptide computed from a previously published ensemble^27^. Lindemann parameters calculated for the backbone **(A)** and side chains **(B)** with the liquid-solid transition and residue-mean indicated. Error bars indicate the 95th percentile of the mean of a bootstrap sample over all 5119 trajectories in the ensemble.

